# HiCBin: Binning metagenomic contigs and recovering metagenome-assembled genomes using Hi-C contact maps

**DOI:** 10.1101/2021.03.22.436521

**Authors:** Yuxuan Du, Fengzhu Sun

## Abstract

Recovering high-quality metagenome-assembled genomes (MAGs) from complex microbial ecosystems remains challenging. Conventional shotgun-based binning approaches may encounter barriers when multiple samples are scarce. Recently, high-throughput chromosome conformation capture (Hi-C) has been applied to simultaneously study multiple genomes in natural microbial communities. Several Hi-C-based binning pipelines have been put forward and yielded state-of-the-art results using a single sample. We conclude that normalization and clustering are two vital steps in the Hi-C-based binning analyses, and develop HiCBin, a novel open-source pipeline, to resolve high-quality MAGs utilizing Hi-C contact maps. HiCBin employs the HiCzin normalization method and the Leiden community detection algorithm based on the Potts spin-glass model and includes the spurious contact detection into binning pipelines for the first time. Using the metagenomic yeast sample with a perfect ground truth of contigs’ species identity, we comprehensively evaluate the impacts on the binning performance of different normalization methods and clustering algorithms from the HiCBin and other available metagenomic Hi-C analysis pipelines, demonstrate that the HiCzin and the Leiden algorithm achieve the best binning accuracy, and show that the spurious contact detection can improve the retrieval performance. We also validate our method and compare the capability to recover high-quality MAGs of HiCBin against other state-of-the-art Hi-C-based binning tools including ProxiMeta, bin3C, and MetaTOR, and one popular shotgun-based binning software MetaBAT2 on a human gut sample and a wastewater sample. HiCBin provides the best performance and applicability in resolving MAGs and is available at https://github.com/dyxstat/HiCBin.

## 1 Introduction

Microbial communities consist of a wide range of microorganisms with many unexploited enzymes and metabolic potentials encoded in the genomes of these diverse species [18, 45]. Traditional pure cultures grown in the laboratory are insufficient to explore the microdiversity because most of microorganisms cannot be cultivated [43, 47]. As a culture-independent genomic approach, metagenomics avoids the isolation or cultivation of microorganisms and provides a broad aspect of the community structure and the functional capabilities present in complex ecosystems [17, 48].

The employment of next-generation sequencing techniques revolutionized metagenomic studies. Genomic fragments are directly sequenced from the microbial ecosystems, generating tremendous read libraries from various environments, such as the human gut, soil, and ocean water [32]. However, the information on the genome identity of reads is lost due to the randomness of the whole-genome shotgun sequencing (WGS) [7]. To retrieve such information, metagenomic analysis assembles the WGS reads into relatively longer contigs. Then, assembled contigs are clustered into metagenome-assembled genomes (MAGs) [19]. This process is usually referred to as metagenomic binning. Traditional shotgun-based binning approaches to the accurate retrieval of MAGs depend on the contig similarity measurements from GC-content, tetra-mer composition, and/or coabundance feature of the contigs across multiple samples [1, 21, 22, 33, 52]. Although experiments have demonstrated that co-abundance profiles across a series of samples enable the discovery of new microbial organisms effectively [37], the requirement for enough samples to obtain reliable coabundance relationships between contigs may not be satisfied due to the cost restriction and limited capability to collect samples, impairing the effectiveness of these conventional binning approaches.

High-throughput chromosome conformation capture (Hi-C) [31] is a DNA proximity technique displaying a great potential to break through the barriers in the metagenomic binning domain. Hi-C technique generates millions of paired-end reads linking DNA fragments in close proximity within cells and has already been utilized to explore topologically associated domains and the compartment property of the mammalian genomes [10, 31]. When applied to metagenomics (metagenomic Hi-C), Hi-C technique is combined with traditional shotgun sequencing and has shown great capabilities to the genome binning and the simultaneous retrieval of high-quality MAGs from a single sample [3, 5].

Metagenomic Hi-C-based binning analysis usually adheres to a standard procedure. Short reads are generated by shotgun sequencing from the microbial community sample. In parallel, paired-end Illumina Hi-C sequence reads are obtained from the same sample. Contigs are assembled from the shotgun sequencing reads and paired-end Hi-C reads are mapped to the assembled contigs to generate raw contact maps, which are then normalized to correct the strong experimental biases. Finally, normalized contact maps are clustered to construct draft genomic bins. From this standard procedure, we can conclude that there are two vital steps directly influencing the binning performance: normalization and clustering. Indeed, several Hi-C-based binning pipelines have been developed using different strategies to do normalization and clustering. ProxiMeta [39], a commercial metagenomic genome binning platform, took the abundance of the contigs into account to normalize the raw Hi-C contacts and then clustered contigs into genome bins using a proprietary MCMC-based algorithm based on their Hi-C linkages [39]. MetaTOR [2] and bin3C [9] are two state-of-the-art open-source pipelines. MetaTOR divided the raw Hi-C contacts by the product of coverage of contig pairs and then applied the Louvain algorithm with the classical Newman-Girvan criterion to cluster contigs [2]. Besides the Hi-C data, MetaTOR can also process the meta3C datasets [35]. bin3C designed a two-stage normalization method to process the raw contact maps [9]. It first divided raw Hi-C counts by the product of the number of restriction sites, and then used the Knight-Ruiz algorithm [24] to construct a general doubly stochastic matrix. For the clustering step, bin3C utilized Infomap software (v0.19.25) [42] to bin the whole contigs. But comprehensive evaluations of different normalization methods and clustering algorithms for the metagenomic Hi-C binning remain sparse [8].

Recently, new research put forward three explicit experimental biases for raw metagenomic Hi-C contacts: the number of restriction sites, contig length, and contig coverage, and has demonstrated that normalization methods in publicly available Hi-C-based binning pipelines cannot correct all explicit biases [11]. Those pipelines implementing normalization by the number of restriction sites cannot even be carried out when the restriction enzymes employed in Hi-C experiments are not specified. Moreover, spurious inter-species contacts (Hi-C contacts between contig pairs from different species) derived from the ligation of DNA fragments between closely related species weakened the interpretability of the Hi-C data [46]. However, none of available Hi-C-based binning pipelines attempted to detect and remove the spurious contacts. As for the clustering step, though several community detection algorithms have been employed to cluster the contigs, those clustering algorithms were not sufficiently effective in general applications [8].

To solve these problems, we develop HiCBin, a new open-source metagenomic Hi-C-based binning pipeline, to recover high-quality MAGs. We employ HiCzin, a novel normalization method designed for metagenomic Hi-C contact maps, to remove the experimental biases. HiCBin also detects and removes the spurious inter-species contacts for the first time among all Hi-C-based binning pipelines. In the clustering step, the Leiden algorithm [49], an advanced modularity-based community detection algorithm, is introduced to the metagenomic binning domain. The Leiden algorithm has proved to be strongly preferable to one of the most popular community detection algorithms, the Louvain algorithm in the experimental benchmarks [4, 49]. We also select a general and flexible modularity function based on the Reichardt and Bornholdt’s Potts model to overcome the resolution limit of the classical Newman-Girvan criterion utilized in the MetaTOR [14, 41]. Using a synthetic metagenomic sample with a perfect ground truth of all contigs [5], we evaluated the retrieval performance of available normalization methods and clustering algorithms from the HiCBin and other Hi-C-based analysis pipelines, and assessed the impacts of spurious contact detection step on the binning results. Finally, we compared the MAG retrieval ability of HiCBin against all the other state-of-the-art Hi-C-based binning pipelines: ProxiMeta, bin3C, and MetaTOR [2, 9, 39], and one widely used shotgun-based binning software MetaBAT2 [23] on a human gut dataset [39] and a wastewater dataset [46].

## 2 Materials and Methods

### 2.1 Datasets

We analyzed the performance of our genome binning tool HiCBin on three published metagenomic Hi-C datasets [5, 39, 46].

#### 2.1.1 Synthetic metagenomic yeast community

The synthetic metagenomic yeast (M-Y) sample consist of 16 yeast strains from 13 yeast species (BioProject: PRJNA245328, Accession: SRR1263009 and SRR1262938) [5]. Nextera DNA Sample Preparation Kit (Illumina) was employed to prepare the shotgun library (SRR1262938). Hi-C library (SRR1263009) was created using HindIII and NcoI restriction endonuclease (NEB). Pairedend sequencing of Hi-C reads was performed using the HiSeq and MiSeq Illumina platforms. Raw WGS dataset contains 85.7 million read pairs (101 bp per read) and raw Hi-C dataset includes 81 million read pairs (100 bp per read). As the reference genomes of all strains in the metagenomic sample are known, we can determine the true species identity of the assembled contigs by aligning assembled contigs to reference genomes at the species level and then construct a gold standard to validate the whole-community genome binning performance (described below).

#### 2.1.2 Real microbiome communities from a human gut sample and a wastewater sample

To compare HiCBin to a proprietary metagenome genome binning service (ProxiMeta), two publicly available metagenomic Hi-C sequencing datasets constructed by the ProxiMeta Hi-C kit (Phase Genomics, Seattle, WA, USA) were chosen [39, 46].

The first dataset was derived from a fecal sample of a human subject (BioProject: PRJNA413092, Accession: SRR6131122, SRR6131123, and SRR6131124) [39]. Two four-cutter restriction enzymes MluCI and Sau3AI (New England Biolabs) were utilized to construct two different Hi-C libraries (SRR6131122, SRR6131124). The shotgun and Hi-C libraries were sequenced by a Illumina HiSeqX platform at 151 bp. The raw shotgun library (SRR6131123) consisted of 250,884,672 read pairs while the sequencing of two Hi-C libraries produced 41,733,770 read pairs (Sau3AI library) and 48,798,091 read pairs (MluCI library), respectively.

The second dataset was generated from a wastewater (WW) sample (BioProject: PRJNA506462, Accession: SRR8239392 and SRR8239393) [46]. The shotgun library (SRR8239393) and Hi-C library (SRR8239392) were prepared using the DNeasy PowerWater kit and ProxiMeta Hi-C kit, respectively. After sequencing both libraries by HiSeq 4000 at 150 bp, 269,312,499 and 95,284,717 read pairs for the wastewater shotgun and Hi-C libraries were produced, respectively. Noticeably, the restriction enzymes employed in the experiment were unspecified due to the proprietary Hi-C preparation kit.

### 2.2 Preprocessing raw reads

A standard cleaning procedure was applied to all raw WGS and Hi-C read libraries using bbduk from the BBTools suite (v37.25) [6]. We discarded short reads below 50 bp at each cleaning step. Adaptor sequences were removed by bbduk with parameter ‘ktrim=r k=23 mink=11 hdist=1 minlen=50 tpe tbo’ and reads were quality-trimmed using bbduk with parameters ‘trimq=10 qtrim=r ftm=5 minlen=50’. Then, the first 10 nucleotides of each read were trimmed by bbduk with parameter ‘ftl=10’.

### 2.3 Shotgun assembly

De novo metagenome assembly was produced by MEGAHIT (v1.2.9) [28] with parameters ‘-mincontig-len 300 -k-min 21 -k-max 141 -k-step 12 -merge-level 20,0.95’ using processed shotgun reads and contigs below 1 kb were discarded (Table 1).

**Table 1:** Statistics of assembled contigs for three datasets.

| Dataset | Contigs ≥ 1 kbp | N50 | Average length | Total length |
| --- | --- | --- | --- | --- |
| Metagenomic yeast | 6,566 | 63,305 | 10,781 | 126,030,343 |
| Human gut | 105,267 | 14,166 | 5044 | 530,969,816 |
| Wastewater | 752,580 | 2977 | 2538 | 1,910,562,642 |

### 2.4 Hi-C read alignment

For the Hi-C library, only paired reads were retained for the downstream workflow. All identical PCR optical and tile-edge duplicates for Hi-C paired-end reads were removed by the script ‘clumpify.sh’ from BBTools suite (v37.25) [6] with default parameters. Then, processed Hi-C paired-end reads were aligned to assembled contigs using BWA MEM (v0.7.17) [29] with parameter ‘-5SP’. Samtools (v1.9) [30] with parameters ‘view -F 0×904’ were subsequently applied to the resulting BAM files to remove unmapped reads (0×4), secondary alignments (0×100), and supplementary alignments (0×800). Alignments with low quality (nucleotide match length <30 or mapping score <30) were filtered out.

### 2.5 Generating Hi-C contact maps

Raw contig-contig interactions were aggregated as contacts by counting the number of alignments bridging two contigs [25]. As contacts reflect the proximity extent between contigs, only pairs of Hi-C reads aligned to different contigs were retained to generate the contact maps. We then denote the contig signal as the number of Hi-C reads mapped to the contig. In the Hi-C experiment, shorter contigs with smaller signals tend to have much higher variance, weakening the normalization and clustering stability in the downstream analysis. To get rid of these deleterious impacts, restrictions on minimum contig length (default, 1000 bp) and minimum contig signal (default, two) were imposed to filter problematic contigs. We discarded contigs failing to satisfy either of the two limitations. In this way, raw contact maps were generated from the Hi-C read pairs to measure the interactions between contigs.

### 2.6 Normalizing the raw contact map

Apart from chromosomal contacts of interest, two types of experimental biases have been reported to have substantial effects on raw metagenomic Hi-C contact maps, rendering the normalization of Hi-C contact maps essential for the binning [11]. Therefore, we employed HiCzin, a state-of-the-art metagenomic Hi-C normalization method, to remove the influences of biases [11]. We denote the intra-species contacts as the count of Hi-C interactions between contigs from the same species and refer to the non-zero intra-species contacts as valid contacts. Based on the assumption that the observed intra-species contacts follow zero-inflated negative binomial distribution, the HiCzin model corrected biases of the number of restriction sites, length and coverage of contigs while taking the biases of unobserved Hi-C interactions into account. Considering the restriction enzymes utilized in Hi-C experiments are unspecified in some real situations, HiCzin also developed a special mode (HiCzin LC) that merely normalized raw Hi-C contact maps by the length and coverage of contigs. The main procedure of the HiCzin normalization contained the following steps:

#### 2.6.1 Computing the coverage of assembled contigs

The coverage of contigs was computed using MetaBAT2 (v2.12.1) [22, 23] script: ‘jgi summarize bam contig depths’.

#### 2.6.2 Generating the intra-species contacts

To generate the observations of the intra-species contacts, TAXAassign (v0.4) ^1^ [20] was utilized to resolve the taxonomic assignment of contigs using NCBI’s Taxonomy and its nt database with parameters ‘-p -c 20 -r 10 -m 98 - q 98 -t 95 -a “60,70,80,95,95,98” -f’. We discarded indeterministic assignment results with ‘unclassified’ labels at the species level. In this way, some contigs could be unambiguously annotated at the species level. Intra-species pairs were subsequently constructed by pairwise combining contigs assigned to the same species and corresponding contacts were regarded as the samples of the intra-species contacts.

#### 2.6.3 Fitting the regression model

All intra-species contact samples were utilized to fit the HiCzin or HiCzin LC model, combining the negative binomial distribution of the intra-species contacts with a mass distribution of unobserved contacts. The residues of the counting part served as the normalized contacts.

### 2.7 Removing spurious contacts

Based on the expectation that the magnitude of the normalized spurious contacts by HiCzin or HiCzin LC to be significantly smaller than that of the normalized valid contacts [11], we discarded the normalized contacts below a threshold as spurious contacts, and determined the threshold such that less than a preselected percentage (default, 5%) of non-zero intra-species contact samples generated by TAXAassign are incorrectly identified as spurious contacts. This percentage reflected the acceptable fraction of losses of the valid contacts in the whole data. In this way, we could keep the proportion of incorrectly discarded intra-species contacts under control while discarding most of the spurious contacts as shown in previous work [11].

### 2.8 Clustering by the Leiden algorithm

Compared to classical clustering algorithms [53], such as K-means, K-medoids and Gaussian mixture model, community detection algorithms do not require the input of the number of clusters [27]. The Louvain algorithm is one of the most popular community detection algorithms used to cluster contigs based on metagenomic Hi-C data [2, 4, 34]. As a hierarchical agglomerative method, the Louvain algorithm takes a two-stage greedy approach to optimizing the modularity function [15]. Specifically, the algorithm iterates and assigns each node to a community such that the local movement will increase the modularity function, followed by aggregating the network. This process repeats iteratively until convergence. However, many recent experiments have shown that such kind of local movement may identify disconnected communities within groups [49], weakening the interpretability of the MAG retrieval. To address this problem, we introduced the Leiden algorithm [49], one of the most novel and advanced community detection algorithms, into the metagenomic Hi-C binning domain. The Leiden algorithm is also a modularity-based algorithm and improves the Louvain algorithm by refining the partition before aggregating the network to guarantee the connectivity of each community. Additionally, the Leiden algorithm is much faster than the Louvain algorithm by a fast local move approach [49]. In practice, the Leiden algorithm has achieved good performance in clustering transcriptomics data [16] and gene expression data [51]. Thus, we applied this state-of-the-art clustering algorithm to the normalized Hi-C contact maps.

Selecting the objective function is crucial for modularity-based algorithms. The classical Newman-Girvan criterion [15] is restricted by a resolution limit and may fail to identify small communities [14]. To overcome the resolution limit in the modularity context, we selected a general and flexible measure of community structure based on the Reichardt and Bornholdt’s Potts model [41] as the modularity function of the Leiden algorithm, i.e,

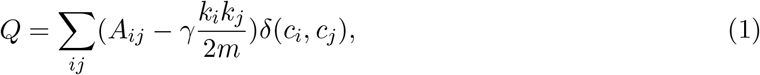

where *Q* is the modularity function, *A* is the adjacency matrix, *k*_*i*_ is the weighted degree of contig *i, m* is the total edge weight, *γ>*0 is a resolution parameter, *c*_*i*_ denotes the community of contig *i* and *δ*(*c*_*i*_, *c*_*j*_) = 1 if *c*_*i*_ = *c*_*j*_ and 0 otherwise. The resolution parameter *γ* in front of the configuration null part controls the relative importance between links within the communities and the null model. Noticeably, this hyperparameter determines the number of communities. Higher resolutions always lead to more communities. Therefore, tuning this parameter is important for the clustering algorithm.

As some contigs have been unambiguously annotated by TAXAassign in the normalization step, we took advantage of these taxonomic assignment results to select an optimal resolution parameter. For each candidate resolution parameter chosen from 1, 5, 10, 15, 20, 25, and 30, we clustered the whole set of contigs and then computed two effective clustering evaluation measures (Appendix): Adjusted Rand Index (ARI) and Normalized Mutual Information (NMI) utilizing those contigs that could be labeled by TAXAassign. We calculated the resolution score as the average of these two measures, and selected the candidate producing the highest resolution score as the optimal value of the hyperparameter.

After determining the resolution parameter, we could finally cluster the whole set of contigs. We set the minimum bin size as default 150 kbp, which was slightly smaller than the minimal length of known bacterial genomes [36]. Only contig bins above 150 kbp were regarded as valid genome bins for the downstream workflow.

### 2.9 Gold standards to evaluate binning performance for the M-Y sample

To evaluate the contig binning results for the metagenomic yeast sample, a perfect ground truth for the species identity of all assembled contigs was constructed. We first downloaded the reference genomes of all 16 yeast strains in the metagenomic sample (Appendix Table A1) [5]. As the analyses were made at the species level, genomes of four strains (FY, CEN.PK, RM11-1A and SK1) from the same species (S. cerevisiae) were merged into one reference genome. Then, assembled contigs were aligned to those 13 reference genomes of all known species by BLASTN [54] with parameters: ‘-perc identity 95 -evalue 1e-30 -word size 50’. The true species identity of the assembled contigs could be determined if there existed any alignment of the contigs to the species’ reference genome (Figure 1). After we obtained the true labels of all contigs, we explored three comprehensive clustering performance metrics (Appendix): Fowlkes-Mallows scores (F-scores), Adjusted Rand Index (ARI), and Normalized Mutual Information (NMI). These three metrics served as the gold standards to evaluate binning performance for the M-Y sample.

**Figure 1:**
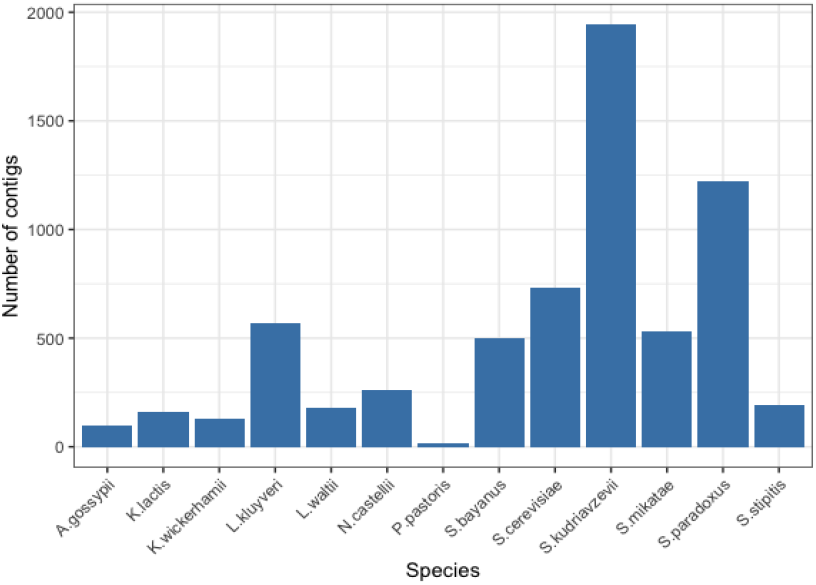
The number of assembled contigs for each of the 13 species in the metagenomic yeast sample.

### 2.10 Analyzing genome bins for the human and wastewater samples

For the human gut and wastewater datasets, as we did not know the ground truth of contig identities, we employed CheckM [38] (v1.1.3, parameter: lineage wf), which searched the single marker genes in the bins, to evaluate draft assembled genomes. According to the CheckM criteria for completeness [39], draft microbial genomes could be assigned to three ranks, i.e., near-complete (≥ 90% completeness, ≤ 10% contamination), substantially complete (≥ 70% and <90% completeness, ≤ 10% contamination), and moderately complete (≥ 50% and <70% completeness, ≤ 10% contamination).

### 2.11 Post-processing on partially contaminated bins

We defined genome bins with completeness larger than 50% and contamination larger than 10% as partially contaminated bins. A post-processing step was designed to clean these partially contaminated bins by re-clustering contigs within each contaminated bin using the Leiden algorithm. As the number of groups within each bin was expected to be small, the resolution parameter was kept to be 1 in the post-processing step. By this means, groups of relatively smaller bins, denoted by sub bins, could be obtained and those sub bins satisfying the minimum bin size requirement were retained and subsequently evaluated for their quality by CheckM.

### 2.12 Comparison to other metagenomic binning pipelines

We compared HiCBin to all the other Hi-C-based metagenome deconvolution pipelines, i.e, Prox-iMeta [39], MetaTOR (v0.1.7) [2], and bin3C (v0.1.1) [9]. We also compared HiCBin with Mata-BAT2 (v2.12.1) [23], which was a conventional binning pipeline using shotgun libraries only and achieved one of the best binning performance in the CAMI Challenge Datasets [44]. MetaTOR, bin3C, and MetaBAT2 are three open-source tools. MetaTOR was implemented with default parameters, followed by the recursive Louvain clustering [2]. bin3C and MetaBAT2 were run with default parameters. As ProxiMeta is a proprietary metagenomic genome binning platform without an open-source pipeline, we reanalyzed two datasets processed by ProxiMeta kit (i.e., the human gut and the wastewater datasets), and compared the CheckM validations of HiCBin to the results of ProxiMeta provided in their supplementary data. We also compared the quality of retrieved genome bins of HiCBin and the other three open software tools on the human gut sample according to the CheckM standard. For the wastewater sample, as the restriction enzymes employed in the proprietary preparation kit were unspecified in the experiment, bin3C could not compute the number of restriction sites on contigs and normalize the raw contacts, resulting in the inapplicability of bin3C. Therefore, we additionally compared the CheckM results of HiCBin to the results of MetaTOR and MetaBAT2.

## 3 Result

### 3.1 Analyses of the metagenomic yeast sample

A total of 6566 contigs longer than 1000 bp were assembled with the total length of 126,030,343 bp (Table 1). There were 4,700,202 Hi-C read pairs subsequently aligned to different contigs. Among the contigs, 5283 of them passed the filtering criteria and represented 80.5% of all assembled contigs and 98.6% of the total length of the entire shotgun assembly. For the normalization step, 2700 contigs were labeled by TAXAassign, generating 847,109 intra-species contacts to fit the HiCzin model. We discarded normalized contacts below 0.178 as spurious contacts. Thereafter, 14 valid genome bins with the bin size larger than 150 kbp were obtained by the Leiden algorithm with a resolution parameter as 1. These valid bins contained 5217 contigs with the total length of 124,120,326 bp, representing 98.5% of the total length of all shotgun assembled contigs longer than 1000 bp. The F-score, ARI, and NMI were 0.905, 0.891, and 0.887, respectively. We then investigated the influences on the binning performance of available normalization methods and clustering algorithms from the HiCBin and other metagenomic Hi-C analysis pipelines and evaluated the impacts of spurious contact detection step on the binning results based on our gold standards.

#### 3.1.1 Exploring the impacts of different normalization methods on binning

Apart from a state-of-the-art normalization method HiCzin [11], several relatively simple metagenomic Hi-C normalization methods have been developed. Beitel et al. [3] divided raw Hi-C contacts by the product of the length of two contigs. MetaTOR [2] normalized raw Hi-C interactions by the geometric mean of the contigs’ coverage. Metaphase [5] and bin3C [9] divided raw Hi-C counts by the product of the number of restriction sites and bin3C used the Knight-Ruiz algorithm [24] to construct a general doubly stochastic matrix after the first step correction. For convenience, we denote those normalization methods by site, length, and coverage as Naive Site, Naive Length, and Naive Coverage, and denote the two-stage normalization method in bin3C as bin3C Norm. We normalized the raw Hi-C contacts by different normalization methods and applied the Leiden algorithm with the same resolution parameter to cluster the contigs. The gold standards were utilized to evaluate the binning results. As shown in Table 2, HiCzin achieved the best binning performance among all five normalization methods.

**Table 2:** Results of different normalization methods followed by the Leiden clustering algorithm.

|  | # of groups | # of contigs | F-score | ARI | NMI |
| --- | --- | --- | --- | --- | --- |
| Naive Site | 13 | 5191 | 0.870 | 0.851 | 0.859 |
| Naive Length | 13 | 5210 | 0.887 | 0.871 | 0.864 |
| Naive Coverage | 11 | 5240 | 0.757 | 0.697 | 0.838 |
| bin3C_Norm | 13 | 5155 | 0.889 | 0.873 | 0.878 |
| HiCzin (HiCBin) | 14 | 5217 | **0.905** | **0.891** | **0.887** |

Note: the optimal values of the results are in **bold**. # of groups represents the number of valid genome bins and # of contigs is the number of contigs in the valid bins.

#### 3.1.2 Comparing different community detection algorithms

Besides the Louvain algorithm and the Leiden algorithm (see Materials and Methods), there are several widely-used network clustering algorithms, such as Markov clustering algorithm [50], Infomap algorithm [42], and Label Propagation algorithm [40]. Markov clustering and Infomap algorithm are both based on flow models and have already been employed to cluster contigs using metagenomic Hi-C contact maps [3, 9]. The label propagation algorithm iteratively repeats a process where each node in the graph adopts the most common label amongst its neighbors. Some studies have already compared part of these community detection algorithm in the benchmarking networks [13, 26]. However, none of these comparisons focused on the metagenomic Hi-C data. In addition, as one of the latest network clustering algorithm, the Leiden algorithm was seldom taken into consideration for the comparison. Therefore, we investigated the performance of different community detection algorithms in clustering metagenomic Hi-C contact graph. We applied multiple algorithms in the clustering step of the HiCBin pipeline and evaluated the valid genome bins generated by different clustering algorithms (Table 3). Both flow-based algorithms generated much more communities than the real number of species with poor clustering quality. The label propagation algorithm could not detect all species. Two modularity-based algorithms (i.e, the Louvain algorithm and the Leiden algorithm) obtained better performance than other algorithms. The Leiden algorithm provided the best binning results and had a significantly improvement to the Louvain algorithm.

**Table 3:** Results of different community detection algorithms based on the contacts normalized by HiCzin.

|  | # of groups | # of contigs | F-score | ARI | NMI |
| --- | --- | --- | --- | --- | --- |
| Label Propagation | 10 | 5026 | 0.675 | 0.593 | 0.732 |
| Markov clustering | 37 | 4699 | 0.544 | 0.470 | 0.682 |
| Infomap | 66 | 3614 | 0.764 | 0.720 | 0.764 |
| Louvain | 14 | 5240 | 0.837 | 0.814 | 0.834 |
| Leiden (HiCBin) | 14 | 5217 | **0.905** | **0.891** | **0.887** |

Note: the optimal values of the results are in **bold**. # of groups represents the number of valid genome bins and # of contigs is the number of contigs in the valid bins.

#### 3.1.3 Exploring the influences of spurious contact detection on HiCBin

As spurious contact detection is a novel step in metagenomic Hi-C binning pipelines, we explored how this new step benefits the HiCBin pipeline. By discarding normalized contacts below 0.178, we could remove 46.9% of spurious contacts while only 5.6% of valid contacts were incorrectly discarded, which was close to our preselected percentage of acceptable incorrectly identified valid contacts in the whole data. Thus, a large fraction of spurious contacts were removed while most of the valid contacts were retained. Moreover, the HiCBin pipeline was also run without the step of spurious contact detection, where the F-score, ARI, and NMI were 0.901, 0.887, and 0.884, respectively. These three standards were increased to 0.905, 0.891, and 0.887 after removing spurious contacts. Therefore, spurious contact detection indeed improved the binning performance.

#### 3.1.4 Comparing different binning pipelines

We compared the binning performance of HiCBin to two publicly available metagenomic binning pipelines, MetaBAT2 and bin3C, on the metagenomic yeast dataset (Table 4). MetaBAT2 is a conventional shotgun-based binning pipeline, and bin3C is an open-source solutions to deconvolute the metagenomic Hi-C datasets (see Materials and Methods). Noticeably, as a publicly available Hi-C-based binning tool, MetaTOR splits contigs into ‘chunks’ of nearly fixed size after the assembly, and attempts to bin these chunks instead of the assembled contigs. It is not valid to compare MetaTOR to other pipelines without splitting based on our gold standards due to different clustering objectives. Therefore, we did not include MetaTOR into the comparison for the metagenomic yeast dataset.

**Table 4:**
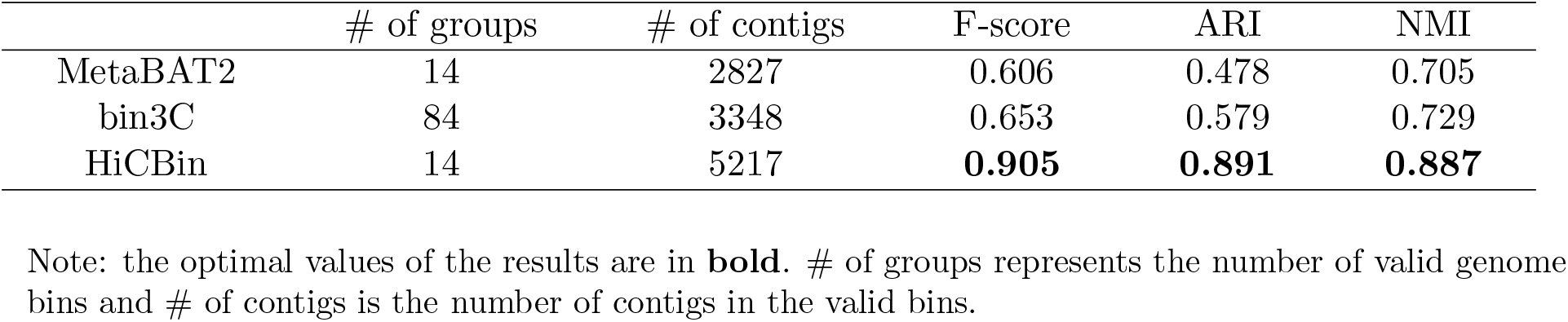
Compare the performance of Hi-C-based binning pipelines: HiCBin and bin3C, and a shotgun-based binning tool MetaBAT2.

|  | # of groups | # of contigs | F-score | ARI | NMI |
| --- | --- | --- | --- | --- | --- |
| MetaBAT2 | 14 | 2827 | 0.606 | 0.478 | 0.705 |
| bin3C | 84 | 3348 | 0.653 | 0.579 | 0.729 |
| HiCBin | 14 | 5217 | **0.905** | **0.891** | **0.887** |

Note: the optimal values of the results are in **bold**. # of groups represents the number of valid genome bins and # of contigs is the number of contigs in the valid bins.

Without using Hi-C information, MetaBAT2 only binned fewer than half of the contigs with relatively poor quality. bin3C had better binning results than MetaBAT2. However, the number of communities generated by bin3C was much larger than the real number of species, which was consistent with the result of Infomap algorithm as bin3C utilized Infomap algorithm to do clustering. HiCBin included almost all contigs in the valid genome bins and achieved much better binning performance than all other comparable binning pipelines on the metagenomic yeast dataset.

### 3.2 Analyses of the human gut sample

We assembled 105,267 contigs longer than 1000 bp with a total length of 530,969,816 bp (Table 1). Two Hi-C libraries were merged and a total of 11,633,561 Hi-C sequencing read pairs mapped mates to different contigs. There were 66,809 contigs totaling 457,763,358 bp in length that passed the contig filtering restriction and accounted for 86.2% of the total length of all assembled contigs. In the normalization step, TAXAassign labeled 6225 contigs and generated 850,289 intra-species contact samples to fit the HiCzin model. Normalized contacts below 0.248 were discarded as spurious contacts. The resolution parameter was tuned to be 20 for clustering. After binning from contact maps, 194 valid genome bins were identified with a total size of 434,186,687 bp. Among these bins, 36 valid bins totaling 180,682,506 bp in size were partially contaminated and were subsequently post-processed, generating 180 sub bins with a total bin size of 148,154,265 bp. Therefore, HiCBin recovered 338 valid genome bins in total. These 338 bins ranging from 150,395 bp to 4,884,489 bp had a total size of 401,658,446 bp and represented 75.6% of the total length of the whole assembled contigs.

We compared the MAG retrieval ability of HiCBin to all the other Hi-C-based binning pipelines (ProxiMeta, bin3C, and MetaTOR) and one conventional shotgun-based binning tool (MetaBAT2) according to the CheckM standard (see Materials and Methods). CheckM validation results of ProxiMeta and bin3C came from the supplementary materials of their papers [9, 39]. HiCBin retrieved 67 near-complete, 33 substantially complete, and 12 moderately complete MAGs (Figure 2a). Near-complete MAGs ranged from 1,509,376 bp to 4,884,489 Mbp, while the substantially complete MAGs ranged from 1,549,630 bp to 2,662,928 bp and moderately complete MAGs ranged from 1,336,581 bp to 2,881,511 bp. In comparison, MetaBAT2 generated 282 bins with 30 near-complete, 25 substantially complete, and 22 moderately complete MAGs (Figure 2b). Genomic bins recovered by MetaTOR suffered serious contaminations without the recursive Louvain clustering. The average contamination of bins with completeness larger than 50% was 80.94%. One potential explanation for the high contamination of draft genomes was that the Louvain algorithm in MetaTOR utilized the Newman-Girvan criterion as the modularity function, which could not identify small genomes due to the resolution limit in the complex Hi-C contact network with a large number of contigs, and those unidentified small genomes were merged into other draft genomes. After the recursive binning procedure, the average contamination of bins with completeness above 50% was decreased to 7.8% and 7 near-complete, 12 substantially complete, and 10 moderately complete MAGs were retrieved by MetaTOR. ProxiMeta created 247 bins with 50 near-complete, 14 substantially complete, and 11 moderately complete MAGs while bin3C constructed 138 bins with 60 near-complete, 24 substantially complete, and 11 moderately complete MAGs. These two Hi-C-based binning pipelines had better binning results than MetaBAT2, demonstrating the high resolution of Hi-C data. HiCBin achieved the best performance in resolving high-quality MAGs on the human gut dataset. Moreover, it was reported in the bin3C paper that contigs totaling 290,643,239 bp and representing 40.4% of the total length of the assembly were included in the bins larger than 50 kbp. Even though HiCBin selected a stricter requirement on minimum bin size, our pipeline still improved the total length of contigs in bins by 38%.

**Figure 2:**
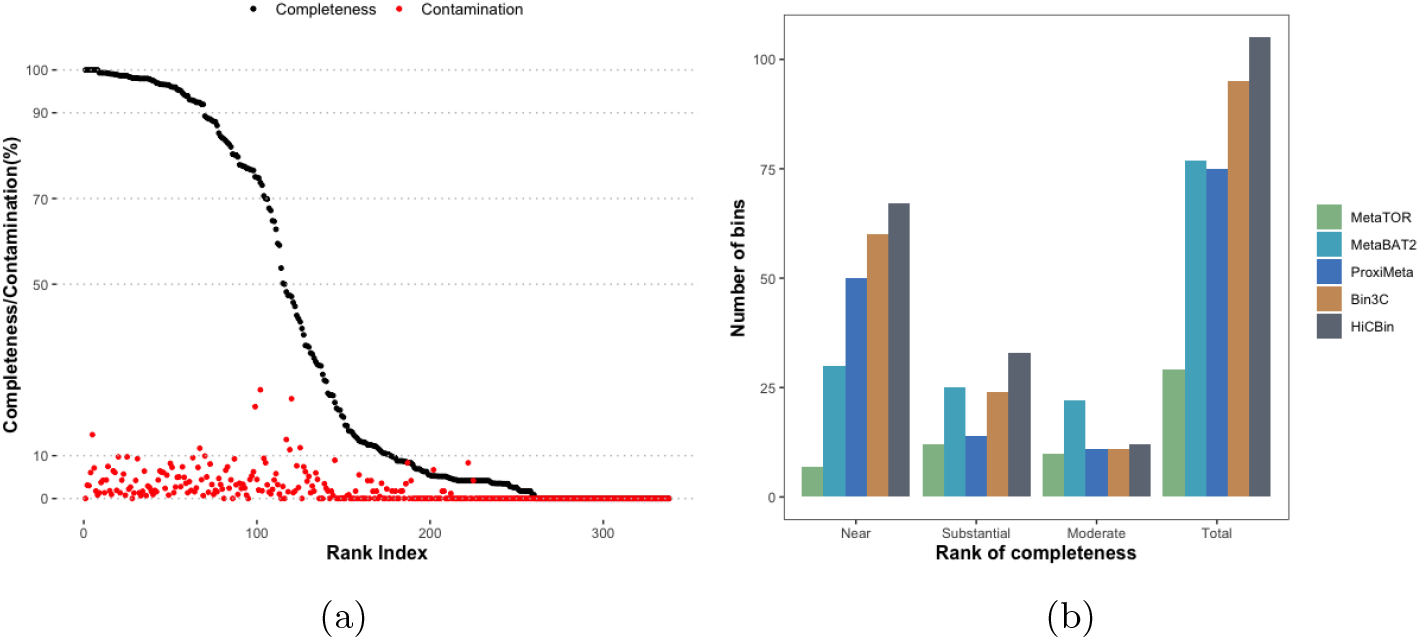
(a) Quality of draft genomic bins retrieved by HiCBin on the human gut dataset; (b) Comparison of different binning pipelines on the human gut dataset according to the CheckM rank for completeness (near-complete: ≥ 90% completeness, ≤ 10% contamination; substantially complete: ≥ 70% and <90% completeness, ≤ 10% contamination; moderately complete: ≥ 50% and <70% completeness, ≤ 10% contamination).

We also explored the impacts of spurious contact detection on this real microbiome community sample. Without spurious contact detection, the HiCBin pipeline could retrieve 63 near-complete and 24 substantially complete MAGs, which were improved to 67 near-complete, 33 substantially complete MAGs after removing spurious contacts, indicating the spurious contact detection step could improve the binning quality.

### 3.3 Analyses of the wastewater dataset

As the wastewater microbiome community sample was processed by the proprietary Hi-C preparation kit, the restriction enzymes utilized in the experiment were not specified, resulting in the lack of the information of the number of restriction sites on contigs. Therefore, HiCzin LC mode were employed for normalization [11]. Moreover, compared to the human gut dataset, the wastewater dataset was much more complicated with 752,580 assembled contigs longer than 1000 bp (Table 1). The total length of the whole assembled contigs was 1,910,562,642 bp. A total of 22,277,042 Hi-C read pairs were aligned to different contigs. There were 493,944 contigs that satisfied the filtering restrictions. These contigs totaled 1,519,539,584 bp in length, representing 79.5% of the total length of the whole assembled contigs longer than 1000 bp. For the normalization step, 9,745,145 intra-species contact samples were generated by TAXAassign to fit the HiCzin LC model and then we discarded normalized contacts below 0.625 as spurious contacts. After tuning the hyper-parameter, contigs were clustered by the Leiden algorithm with resolution parameter as 20. A total of 391 valid genome bins were identified with total size of 1,505,330,057 bp. Among them, 152 bins were partially contaminated. After the post-processing step, 1014 sub bins were generated with the total bin size of 963,363,324 bp. In total, HiCBin reconstructed 1253 valid genome bins with the total size of 1,434,645,324 bp, ranging from 150,008 bp to 8,885,313 bp and accounting for almost 75.1% of the total length of overall shotgun contigs.

According to the CheckM completeness standard, HiCBin retrieved 94 near-complete, 56 substantially complete, and 41 moderately complete MAGs (Figure 3a). Near-complete MAGs ranged from 977,957 bp to 5,330,262 bp, while the substantially complete MAGs ranged from 751,056 bp to 5,316,909 bp and moderately complete MAGs ranged from 780,925 bp to 8,885,313 bp. In comparison to other binning pipelines, HiCBin achieved the best binning results on the wastewater dataset (Figure 3b). Noticeably, bin3C required the input of the names of restriction enzymes as it normalized the raw Hi-C contacts using the number of restriction sites on contigs. Since the enzymes utilized in the experiment were unknown, bin3C was not applicable to this dataset. MetaTOR recovered 11 near-complete, 11 substantially complete, and 7 moderately complete MAGs while MetaBAT2 retrieved 34 near-complete, 60 substantially complete, and 54 moderately complete MAGs. ProxiMeta generated 1288 bins with 15 near-complete, 71 substantially complete, and 107 moderately complete MAGs. Although the total number of MAGs retrieved by ProxiMeta was similar to that of HiCBin, HiCBin generated 94 near-complete MAGs, a gain of 527% against the result of ProxiMeta, indicating that the quality of MAGs retrieved by HiCBin was much higher than ProxiMeta.

**Figure 3:**
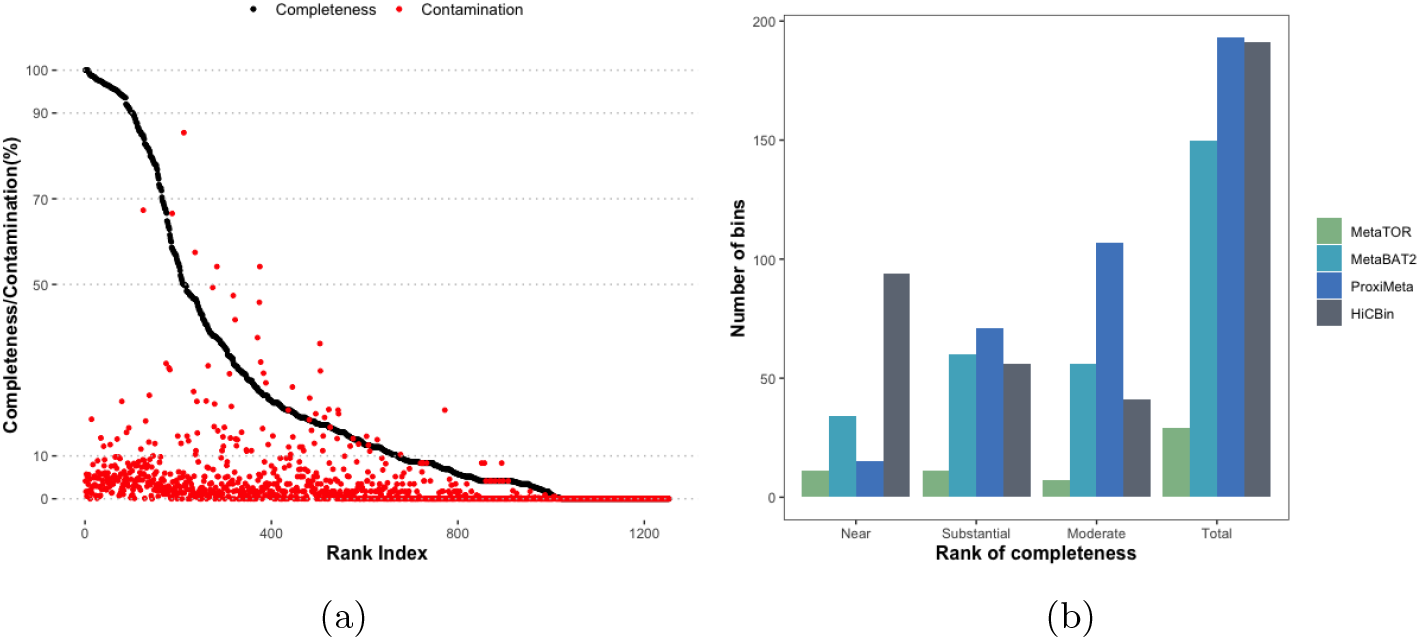
(a) Quality of draft genomic bins retrieved by HiCBin on the wastewater dataset; (b) Comparison of different binning pipelines on the wastewater dataset according to the CheckM rank for completeness (near-complete: ≥ 90% completeness, ≤ 10% contamination; substantially complete: ≥ 70% and <90% completeness, ≤ 10% contamination; moderately complete: ≥ 50% and <70% completeness, ≤ 10% contamination).

### 3.4 Running time of HiCBin, MetaBAT2, and bin3C

HiCBin, bin3C and MetaBAT2 only contains the binning analysis, whereas MetaTOR uses its own workflow and integrates the binning process with standard alignment and annotation softwares. Therefore, it is reasonable to compare running time of binning directly among HiCBin, MetaBAT2, and bin3C. All pipelines were executed on one computing node of 2.40 GHz Intel Xeon Processor E5-2665 provided by the Advanced Research Computing platform at University of Southern California. 50,000 MB memory was allocated to the computing node. The running time of HiCBin and MetaBAT2 on three datasets are shown in Table 5. bin3C was only run on the metagenomic yeast dataset and consumed 21 minutes and 16 seconds, which is slower than HiCBin. Though both Hi-C-based binning pipelines perform better than the shotgun-based binning tool MetaBAT2, MetaBAT2 runs faster than HiCBin and bin3C on the metagenomic yeast dataset. HiCBin consumes less time than MetaBAT2 on the human gut dataset, but consumes more time than MetaBAT2 on the wastewater dataset. This is because it takes relatively long time for HiCBin to generate the observations of the intra-species contacts in the normalization step for the datasets with a huge number of contigs.

**Table 5:**
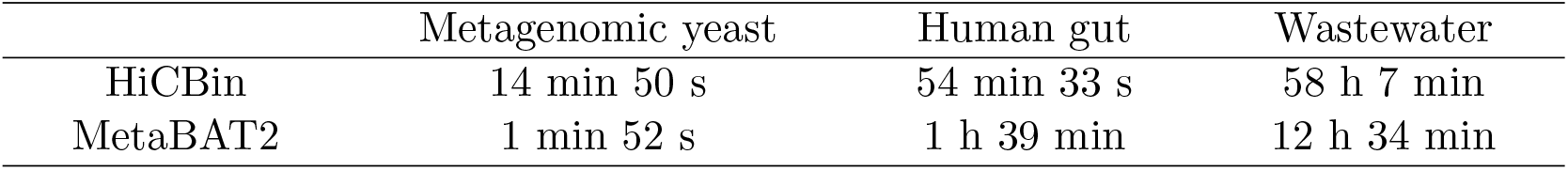
Running time of HiCBin and MetaBAT2 on the yeast, human gut and wastewater datasets..

|  | Metagenomic yeast | Human gut | Wastewater |
| --- | --- | --- | --- |
| HiCBin | 14 min 50 s | 54 min 33 s | 58 h 7 min |
| MetaBAT2 | 1 min 52 s | 1 h 39 min | 12 h 34 min |

## 4 Conclusions and Discussions

We have introduced a new open-source tool HiCBin to resolve high-quality MAGs using Hi-C contact maps. The use of the metagenomic yeast sample with perfect ground truth of contigs’ species identity allowed us to evaluate different normalization methods and clustering algorithms utilized in the HiCBin and other available metagenomic Hi-C analysis pipelines. We found that the HiCzin normalization method and the Leiden algorithm combined with the Potts spin-glass model from the HiCBin provided the best results in genome binning. The spurious contact detection step proved useful in contig binning on both metagenomic yeast dataset and the human gut dataset. Finally, we validated that HiCBin was the best approach to the accurate retrieval of MAGs on the human gut dataset and wastewater dataset compared to all the other Hi-C-based binning pipelines and one traditional shotgun-based software. HiCBin even achieved more than five times improvement against the previous best result in recovering near-complete MAGs for the wastewater dataset. Moreover, HiCBin could include more than 75% of the total length of the whole assembled contigs into bins larger than 150 kbp for both real microbial community samples, representing a marked improvement in the metagenomic binning domain. Notably, the performance of the Hi-C-based binning pipelines are superior to the traditional approach in most instances, indicating the great potential of metagenomic Hi-C data.

As the restriction enzymes utilized in Hi-C experiments are unspecified under some circumstances, those pipelines using the information of the number of restriction sites to do normalization cannot be executed. Employing the HiCzin mode that merely normalized raw Hi-C contact maps by the length and coverage of contigs rendered the HiCBin pipeline more applicable.

From our observation, the selection of minimum bin size is relatively stable. Though different binning pipelines may choose different bin size thresholds, we found that the size of high-quality MAGs determined by CheckM is always much larger than those minimum bin size thresholds.

Although HiCBin performed well in the analysis of Hi-C data, it was not designed for other proximity-ligation techniques such as meta3C [35]. The whole analyses of the metagenomic yeast sample in this paper were at the species level as we found it challenging to annotate contigs at the strain level. In the future, it is better to explore the binning performance with finer resolution. For the two real microbial community datasets, as we don’t know the true species identities of assembled contigs, we employed the CheckM to evaluate the quality of bins. However, the marker-gene based validation may not reflect the true completeness and contamination of recovered MAGs as some contigs don’t contain the marker genes. Therefore, it is worthy of further research on the evaluation of MAG retrieval.

## Appendix

### Evaluation criteria of clustering

#### Fowlkes-Mallows scores

The Fowlkes-Mallows score (FM) is defined as the geometric mean of the pairwise precision and recall, i.e,

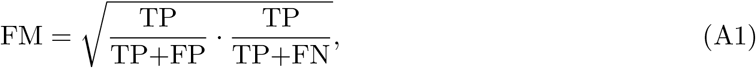

where TP is the number of true positives, FP is the number of false positives, and FN is the number of false negatives.

#### Adjusted Rand Index

Define rand index (RI) as a measure of the percentage of correct decisions made by the clustering algorithm, i.e.,

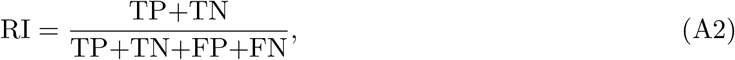

where TP is the number of true positives, TN is the number of true negatives, FP is the number of false positives, and FN is the number of false negatives.

Then, the Adjusted Rand Index (ARI) can be defined as

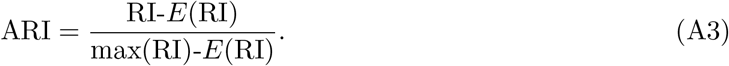

#### Normalized Mutual Information

Let *U* and *V* denote the class labels and cluster labels. Define the entropy of a label set *S* as

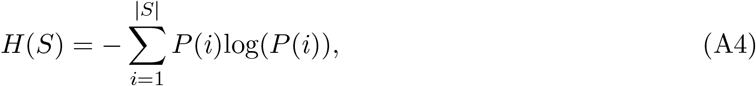

where *P* (*i*) = |*S*_*i*_|*/N* is the probability of an object in class *S*_*i*_.

The mutual information (MI) between *U* and *V* is calculated by:

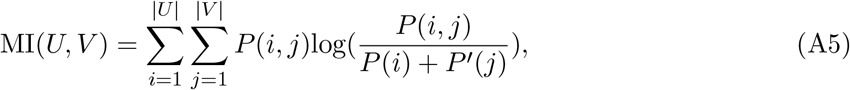

where *P* (*i, j*) = |*U*_*i*_ *∩ V*_*j*_|*/N, P* (*i*) = |*U*_*i*_|*/N*, and *P*′ (*j*) = |*V*_*j*_|*/N*.

Then, the Normalized Mutual Information (NMI) is defined as

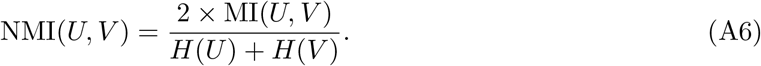

**Table A1:** Reference genomes of the species list in the metagenomic yeast sample.

| Genus | Species | Strain in sample | Reference strain |
| --- | --- | --- | --- |
| Saccharomyces | cerevisiae | FY4H | FY |
| Saccharomyces | cerevisiae | CEN.PK | CEN.PK |
| Saccharomyces | cerevisiae | RM11-1A | RM11-1A |
| Saccharomyces | cerevisiae | SK1 | SK1 |
| Saccharomyces | paradoxus | YDG613 |  |
| Saccharomyces | mikatae | FM356 | IFO 1815 |
| Saccharomyces | kudriavzevii | FM527 | IFO 1802 |
| Saccharomyces | bayanus var. uvarum | YZB5-113 | CBS 7001 |
| Naumovozyma | castellii | 4310 | NRRL Y-12630 |
| Lachancea | waltii | Kwaltii ura3 | NRRL Y-8285 |
| Lachancea | kluyveri | FM628 | CBS 3082 |
| Kluyveromyces | lactis | MW98-8C | NRRL Y-1140 |
| Kluyveromyces | wickerhamii | Y-8286 | UCD 54-210 |
| Ashbya | gossypii | WT | ATCC 10895 |
| Scheffersomyces | stipitis | Y-11545 | CBS 6054 |
| Pichia | pastoris | JC308 | GS115 |

## Declarations

## Abbreviations

Hi-C: high-throughput chromosome conformation capture
MAG: metagenome-assembled genome
WGS: whole-genome shotgun
F-score: Fowlkes-Mallows score
ARI: Adjusted Rand Index
NMI: Normalized Mutual Information

## Acknowledgements

The authors appreciate Ms. Ziye Wang from Fudan University for her kind help with processing the raw data.

## Authors’ contributions

YD and FS conceived the ideas and designed the study. YD implemented the methods, carried out the computational analyses, and drafted the manuscript. FS modified and finalized the paper.

## Availability of data and materials

All the datasets used in this study are publicly available from the NCBI Sequence Read Archive database (http://www.ncbi.nlm.nih.gov/sra). The synthetic metagenomic yeast sample was downloaded under the accession numbers: shotgun library SRR1263009, Hi-C library SRR1262938 [5]. The human gut sample is available through the accession numbers: shotgun library SRR6131123, Hi-C libraries SRR6131122 and SRR6131124 [39]. The accession numbers of the wastewater sample are SRR8239393 for shotgun library and SRR8239392 for Hi-C library. The HiCBin software is freely available at https://github.com/dyxstat/HiCBin [12] under the GNU General Public License version v3. CheckM validation results of the human gut sample and the wastewater sample for Prox-iMeta are available at https://doi.org/10.1101/198713 [39] and https://doi.org/10.1038/s41396-019-0446-4 [46], respectively. Supporting material of bin3C used in comparison is available at https://doi.org/10.1186/s13059-019-1643-1 [9].

## Ethics approval and consent to participate

Not applicable.

## Consent for publication

All authors have approved the manuscript for submission.

## Footnotes

1 https://github.com/umerijaz/TAXAassign

## References

[1] Alneberg, J., Bjarnason, B.S., De Bruijn, I., Schirmer, M., Quick, J., Ijaz, U.Z., Lahti, L., Loman, N.J., Andersson, A.F., Quince, C.: Binning metagenomic contigs by coverage and composition. Nature methods 11(11), 1144–1146 (2014)

[2] Baudry, L., Foutel-Rodier, T., Thierry, A., Koszul, R., Marbouty, M.: MetaTOR: a computational pipeline to recover high-quality metagenomic bins from mammalian gut proximity-ligation (meta3C) libraries. Frontiers in genetics 10, 753 (2019)

[3] Beitel, C.W., Froenicke, L., Lang, J.M., Korf, I.F., Michelmore, R.W., Eisen, J.A., Darling, A.E.: Strain-and plasmid-level deconvolution of a synthetic metagenome by sequencing proximity ligation products. PeerJ 2, e415 (2014)

[4] Blondel, V.D., Guillaume, J.L., Lambiotte, R., Lefebvre, E.: Fast unfolding of communities in large networks. Journal of statistical mechanics: theory and experiment 2008(10), P10008 (2008)

[5] Burton, J.N., Liachko, I., Dunham, M.J., Shendure, J.: Species-level deconvolution of metagenome assemblies with Hi-C–based contact probability maps. G3: Genes, Genomes, Genetics 4(7), 1339–1346 (2014)

[6] Bushnell, B.: BBMap: a fast, accurate, splice-aware aligner. Tech. rep., Lawrence Berkeley National Lab.(LBNL), Berkeley, CA (United States) (2014)

[7] Chen, K., Pachter, L.: Bioinformatics for whole-genome shotgun sequencing of microbial communities. PLoS Comput Biol 1(2), e24 (2005)

[8] DeMaere, M.Z., Darling, A.E.: Deconvoluting simulated metagenomes: the performance of hard-and soft-clustering algorithms applied to metagenomic chromosome conformation capture (3C). PeerJ 4, e2676 (2016)

[9] DeMaere, M.Z., Darling, A.E.: bin3C: exploiting Hi-C sequencing data to accurately resolve metagenome-assembled genomes. Genome biology 20(1), 1–16 (2019)

[10] Dixon, J.R., Selvaraj, S., Yue, F., Kim, A., Li, Y., Shen, Y., Hu, M., Liu, J.S., Ren, B.: Topological domains in mammalian genomes identified by analysis of chromatin interactions. Nature 485(7398), 376–380 (2012)

[11] Du, Y., Laperriere, S.M., Fuhrman, J., Sun, F.: HiCzin: Normalizing metagenomic Hi-C data and detecting spurious contacts using zero-inflated negative binomial regression. bioRxiv (2021). https://doi.org/10.1101/2021.03.01.433489

[12] Du, Y., Sun, F.: HiCBin: Binning metagenomic contigs and recovering metagenome-assembled genomes using Hi-C contact maps. https://github.com/dyxstat/HiCBin (2021)

[13] Emmons, S., Kobourov, S., Gallant, M., Börner, K.: Analysis of network clustering algorithms and cluster quality metrics at scale. PloS one 11(7), e0159161 (2016)

[14] Fortunato, S., Barthelemy, M.: Resolution limit in community detection. Proceedings of the national academy of sciences 104(1), 36–41 (2007)

[15] Girvan, M., Newman, M.E.: Community structure in social and biological networks. Proceedings of the national academy of sciences 99(12), 7821–7826 (2002)

[16] Hagan, T., Cortese, M., Rouphael, N., Boudreau, C., Linde, C., Maddur, M.S., Das, J., Wang, H., Guthmiller, J., Zheng, N.Y., et al.: Antibiotics-driven gut microbiome perturbation alters immunity to vaccines in humans. Cell 178(6), 1313–1328 (2019)

[17] Handelsman, J.: Metagenomics: application of genomics to uncultured microorganisms. Microbiology and molecular biology reviews 68(4), 669–685 (2004)

[18] Hugenholtz, P., Tyson, G.W.: Metagenomics. Nature 455(7212), 481–483 (2008)

[19] Hugerth, L.W., Larsson, J., Alneberg, J., Lindh, M.V., Legrand, C., Pinhassi, J., Andersson, A.F.: Metagenome-assembled genomes uncover a global brackish microbiome. Genome biology 16(1), 1–18 (2015)

[20] Ijaz, U., Quince, C.: TAXAassign v0. 4. https://github.com/umerijaz/TAXAassign (2013)

[21] Imelfort, M., Parks, D., Woodcroft, B.J., Dennis, P., Hugenholtz, P., Tyson, G.W.: GroopM: an automated tool for the recovery of population genomes from related metagenomes. PeerJ 2, e603 (2014)

[22] Kang, D.D., Froula, J., Egan, R., Wang, Z.: MetaBAT, an efficient tool for accurately reconstructing single genomes from complex microbial communities. PeerJ 3, e1165 (2015)

[23] Kang, D.D., Li, F., Kirton, E., Thomas, A., Egan, R., An, H., Wang, Z.: MetaBAT 2: an adaptive binning algorithm for robust and efficient genome reconstruction from metagenome assemblies. PeerJ 7, e7359 (2019)

[24] Knight, P.A., Ruiz, D.: A fast algorithm for matrix balancing. IMA Journal of Numerical Analysis 33(3), 1029–1047 (2013)

[25] Lajoie, B.R., Dekker, J., Kaplan, N.: The Hitchhiker’s guide to Hi-C analysis: practical guidelines. Methods 72, 65–75 (2015)

[26] Lancichinetti, A., Fortunato, S.: Benchmarks for testing community detection algorithms on directed and weighted graphs with overlapping communities. Physical Review E 80(1), 016118 (2009)

[27] Lancichinetti, A., Fortunato, S.: Community detection algorithms: a comparative analysis. Physical review E 80(5), 056117 (2009)

[28] Li, D., Liu, C.M., Luo, R., Sadakane, K., Lam, T.W.: MEGAHIT: an ultra-fast single-node solution for large and complex metagenomics assembly via succinct de Bruijn graph. Bioinformatics 31(10), 1674–1676 (2015)

[29] Li, H.: Aligning sequence reads, clone sequences and assembly contigs with BWA-MEM. arXiv preprint arXiv:1303.3997 (2013)

[30] Li, H., Handsaker, B., Wysoker, A., Fennell, T., Ruan, J., Homer, N., Marth, G., Abecasis, G., Durbin, R.: The sequence alignment/map format and SAMtools. Bioinformatics 25(16), 2078–2079 (2009)

[31] Lieberman-Aiden, E., Van Berkum, N.L., Williams, L., Imakaev, M., Ragoczy, T., Telling, A., Amit, I., Lajoie, B.R., Sabo, P.J., Dorschner, M.O., et al.: Comprehensive mapping of long-range interactions reveals folding principles of the human genome. science 326(5950), 289–293 (2009)

[32] López-García, P., Moreira, D.: Tracking microbial biodiversity through molecular and genomic ecology. Research in microbiology 159(1), 67–73 (2008)

[33] Lu, Y.Y., Chen, T., Fuhrman, J.A., Sun, F.: COCACOLA: binning metagenomic contigs using sequence COmposition, read CoverAge, CO-alignment and paired-end read LinkAge. Bioinformatics 33(6), 791–798 (2017)

[34] Marbouty, M., Baudry, L., Cournac, A., Koszul, R.: Scaffolding bacterial genomes and probing host-virus interactions in gut microbiome by proximity ligation (chromosome capture) assay. Science advances 3(2), e1602105 (2017)

[35] Marbouty, M., Cournac, A., Flot, J.F., Marie-Nelly, H., Mozziconacci, J., Koszul, R.: Metagenomic chromosome conformation capture (meta3C) unveils the diversity of chromosome organization in microorganisms. Elife 3, e03318 (2014)

[36] Nakabachi, A., Yamashita, A., Toh, H., Ishikawa, H., Dunbar, H.E., Moran, N.A., Hattori, M.: The 160-kilobase genome of the bacterial endosymbiont Carsonella. Science 314(5797), 267–267 (2006)

[37] Nielsen, H.B., Almeida, M., Juncker, A.S., Rasmussen, S., Li, J., Sunagawa, S., Plichta, D.R., Gautier, L., Pedersen, A.G., Le Chatelier, E., et al.: Identification and assembly of genomes and genetic elements in complex metagenomic samples without using reference genomes. Nature biotechnology 32(8), 822–828 (2014)

[38] Parks, D.H., Imelfort, M., Skennerton, C.T., Hugenholtz, P., Tyson, G.W.: CheckM: assessing the quality of microbial genomes recovered from isolates, single cells, and metagenomes. Genome research 25(7), 1043–1055 (2015)

[39] Press, M.O., Wiser, A.H., Kronenberg, Z.N., Langford, K.W., Shakya, M., Lo, C.C., Mueller, K.A., Sullivan, S.T., Chain, P.S., Liachko, I.: Hi-C deconvolution of a human gut microbiome yields high-quality draft genomes and reveals plasmid-genome interactions. biorxiv p. 198713 (2017). https://doi.org/10.1101/198713

[40] Raghavan, U.N., Albert, R., Kumara, S.: Near linear time algorithm to detect community structures in large-scale networks. Physical review E 76(3), 036106 (2007)

[41] Reichardt, J., Bornholdt, S.: Statistical mechanics of community detection. Physical review E 74(1), 016110 (2006)

[42] Rosvall, M., Bergstrom, C.T.: Maps of random walks on complex networks reveal community structure. Proceedings of the National Academy of Sciences 105(4), 1118–1123 (2008)

[43] Sait, M., Hugenholtz, P., Janssen, P.H.: Cultivation of globally distributed soil bacteria from phylogenetic lineages previously only detected in cultivation-independent surveys. Environmental microbiology 4(11), 654–666 (2002)

[44] Sczyrba, A., Hofmann, P., Belmann, P., Koslicki, D., Janssen, S., Dröge, J., Gregor, I., Majda, S., Fiedler, J., Dahms, E., et al.: Critical assessment of metagenome interpretation—a benchmark of metagenomics software. Nature methods 14(11), 1063–1071 (2017)

[45] Simon, C., Daniel, R.: Metagenomic analyses: past and future trends. Applied and environmental microbiology 77(4), 1153–1161 (2011)

[46] Stalder, T., Press, M.O., Sullivan, S., Liachko, I., Top, E.M.: Linking the resistome and plasmidome to the microbiome. The ISME journal 13(10), 2437–2446 (2019)

[47] Stevenson, B.S., Eichorst, S.A., Wertz, J.T., Schmidt, T.M., Breznak, J.A.: New strategies for cultivation and detection of previously uncultured microbes. Applied and environmental microbiology 70(8), 4748–4755 (2004)

[48] Streit, W.R., Schmitz, R.A.: Metagenomics–the key to the uncultured microbes. Current opinion in microbiology 7(5), 492–498 (2004)

[49] Traag, V.A., Waltman, L., Van Eck, N.J.: From Louvain to Leiden: guaranteeing well-connected communities. Scientific reports 9(1), 1–12 (2019)

[50] Van Dongen, S.M.: Graph clustering by flow simulation. Ph.D. thesis (2000)

[51] Veres, A., Faust, A.L., Bushnell, H.L., Engquist, E.N., Kenty, J.H.R., Harb, G., Poh, Y.C., Sintov, E., Gürtler, M., Pagliuca, F.W., et al.: Charting cellular identity during human in vitro β-cell differentiation. Nature 569(7756), 368–373 (2019)

[52] Wu, Y.W., Tang, Y.H., Tringe, S.G., Simmons, B.A., Singer, S.W.: MaxBin: an automated binning method to recover individual genomes from metagenomes using an expectation-maximization algorithm. Microbiome 2(1), 1–18 (2014)

[53] Xu, R., Wunsch, D.: Survey of clustering algorithms. IEEE Transactions on neural networks 16(3), 645–678 (2005)

[54] Ye, J., McGinnis, S., Madden, T.L.: BLAST: improvements for better sequence analysis. Nucleic acids research 34(suppl_2), W6–W9 (2006)

